## Supplementary material for "Multi-layered ecological interactions determine growth of clinical antibiotic-resistant strains within human microbiomes"

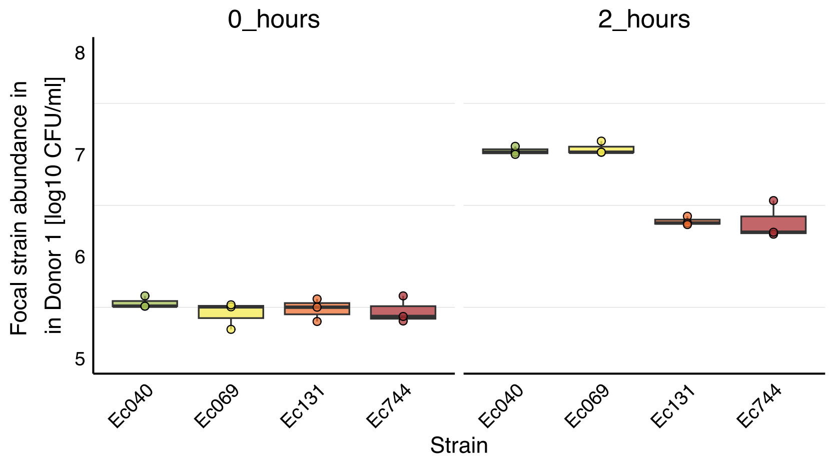


Figure S1. Incoming strains' growth variability in 'live' gut microcosms after two hours. Growth of each introduced resistant strain over two hours in microcosms prepared with the microbiome sample from Donor1. Three replicates are shown for each strain after inoculation at similar densities (0 hours) and after 2 hours. We carried out this experiment to determine whether variable abundances observed in the 'live' microcosm assay (Fig. 1 in the main manuscript) after two hours (the time point at which we introduced the antibiotic treatment, after an initial two-hour reconditioning phase), could be explained by variable growth during the reconditioning phase. The differences among strains after two hours here show a similar pattern as in the 'live' microcosm assay after 2 hours (Fig. 1), suggesting this is possible. Two further lines of evidence suggest strain differences at the end of the 'live' microcosm assay are unlikely to have resulted from variable inoculum densities or early-phase abundances. First, in some treatment groups in the 'live' microcosm assay, the same focal strain reached different final abundances in different treatments, even after having similar abundances after 2 hours (e.g., strain Ec131 with Donor1 vs. Donor2 and Donor3; Fig. 1). Second, if we exclude the strain with the lowest 2h abundance (Ec744), we still detect average differences in final abundance among strains in the absence of antibiotics (strain effect in Fig. 1 tested by two-way ANOVA excluding Ec744 treatments: $F(2,18)= 40$, $p<0.001$), even though the remaining three strains had similar 2h abundances. This suggests there are strain differences in invasion success not driven by variable inoculum size.

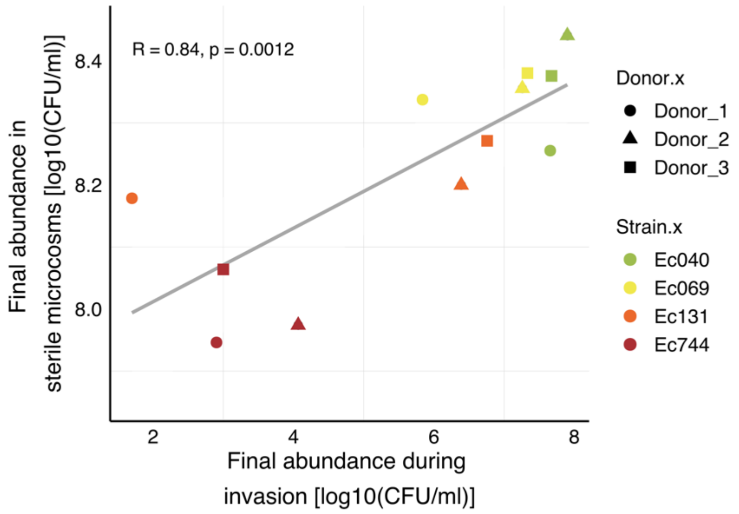


Figure S2. Positive correlation between final abundance of incoming strains after incubation in sterilised microcosms (local abiotic conditions) (Fig. 2) and in 'live' gut microcosms from the 'live' microcosm assay (Fig. 1). Colours indicate the different incoming resistant strains and shapes indicate the different donor samples (see legend). Each point gives the mean of three replicates on both axes.

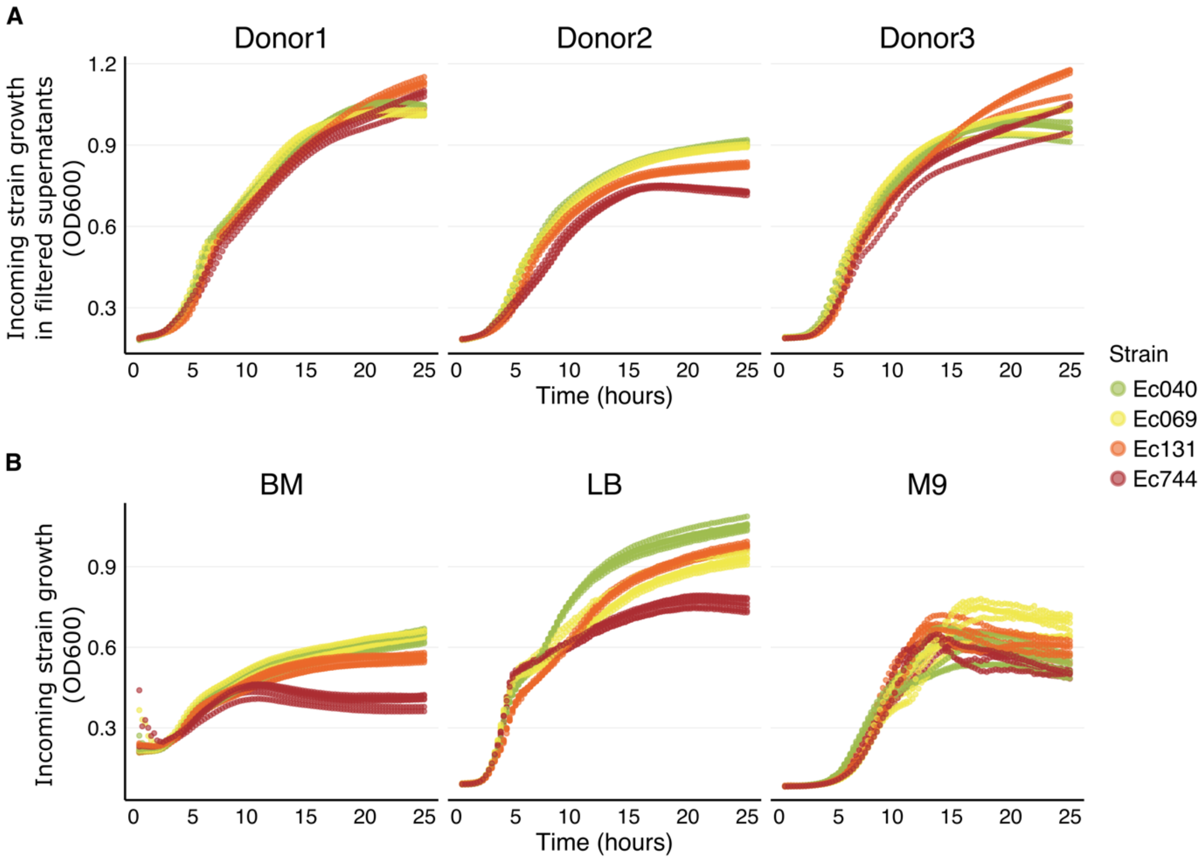


Figure S3. Growth curves of each *E. coli* incoming strain in various conditions. (A) Growth experiments in filtered versions of faecal slurries from each of the same three healthy human donors as in the main experiment (panels left to right). These curves also help to rule out the possibility of phages present in the supernatants having a role on the population growth performance. (B) Growth experiments in three types of sterile media (BM: Basal Medium, added to faecal slurry to prepare microcosms in the main experiment; LB: Lysogeny Broth; M9: M9 minimal salts supplemented with glucose; see Materials and Methods in the main text). All these experiments were performed in microplates in aerobic conditions. Each series of points shows one of four biological replicates in each combination of strain and condition.

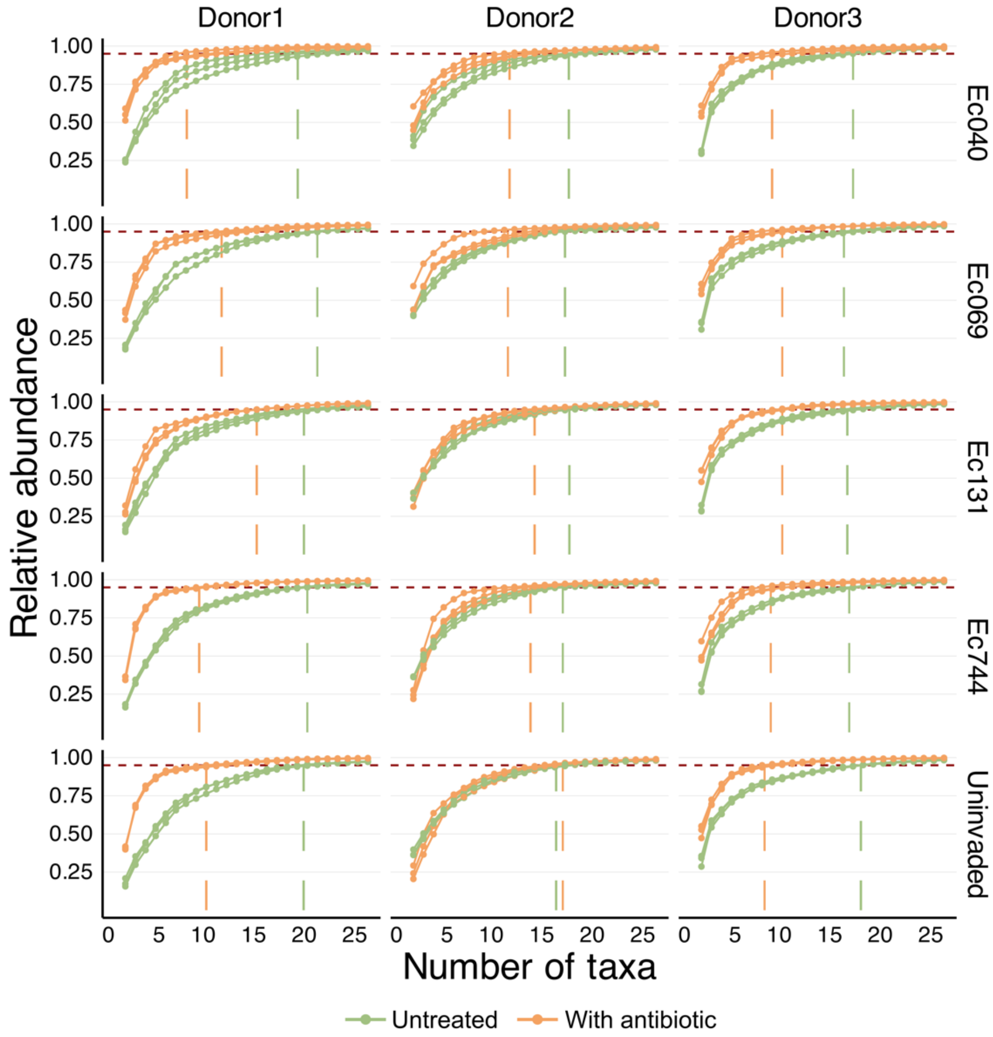


Figure S4. Taxa (Genus) accumulation showing relative abundance after 48 hours in microcosms untreated (green) and treated with antibiotics (orange). The threshold line indicates 95% of the total abundance. The three series in each plot show three replicate microcosms.

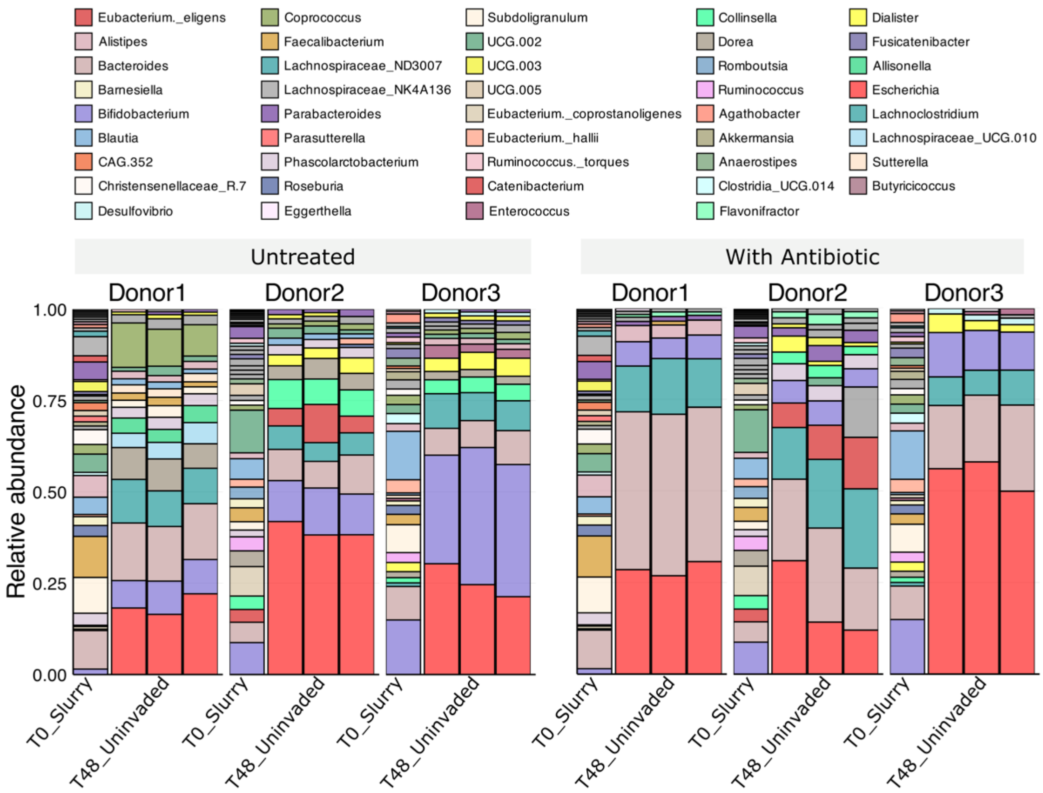


Figure S5. Relative abundance of genera in faecal slurries prior to the microcosm's preparation and to inoculation/incubation (*x*-axis: T0_Slurry) and in microcosms where no focal strain was introduced (uninoculated) after 48 hours of anaerobic incubation (*x*-axis: T48_Uninvaded) for each human donor sample (labelled at top), in the absence and presence of antibiotics (7.2 µg/ml ampicillin) (panel labels above). This plot illustrates the changes in community composition from the initial faecal slurries to the control communities after 48 hours under lab conditions. The original microbial composition of the slurries before the experiment is shown only as a methodological reference; however, the final composition under laboratory conditions is the only one considered for analysis and comparisons in this study. Note that our comparison of relative abundance post-inoculation was made against the uninvaded microcosms after 48 hours, rather than the original slurry composition. In microcosms incubated without an incoming strain and without antibiotics, some resident taxa increased in relative abundance while others declined, although the identities of the top genera (representing the 95% of the total composition) remained similar (12 out of 20 were unchanged over time with Donor1, 12 with Donor2 and 10 with Donor3).

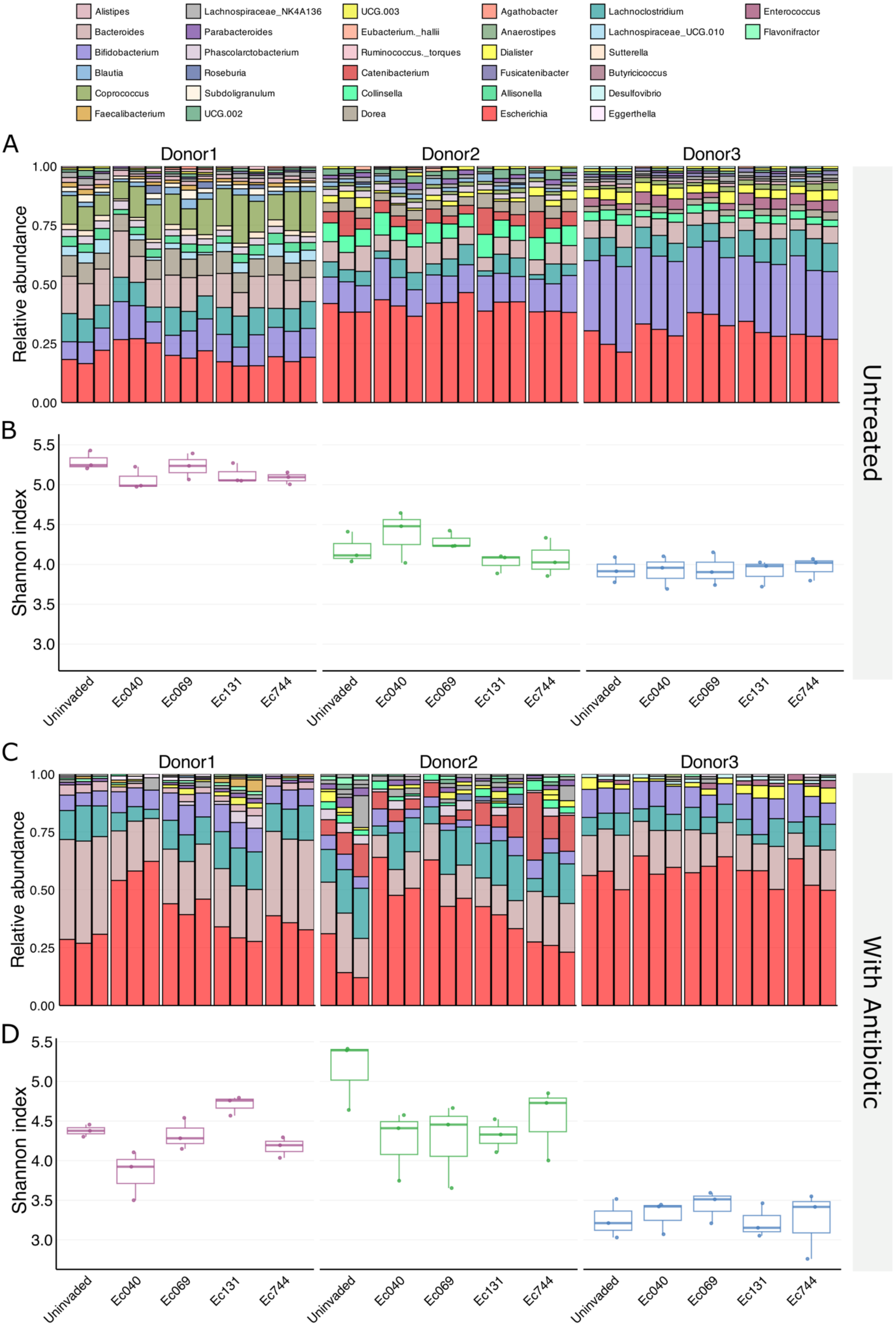

Alistipes
Bacteroides
Bifidobacterium
Blautia
Coprococcus
Faecalibacterium
Lachnospiraceae_NK4A136
Parabacteroides
Phascolarctobacterium
Roseburia
Subdoligranulum
UCG.002
UCG.003
Eubacterium._hallii
Ruminococcus._torques
Catenibacterium
Collinsella
Dorea
Agathobacter
Anaerostipes
Dialister
Fusicatenibacter
Allisonella
Escherichia
Lachnoclostridium
Lachnospiraceae_UCG.010
Sutterella
Butyricicoccus
Desulfovibrio
Eggerthella
Enterococcus
Flavonifractor
A
Donor1
Donor2
Donor3
Relative abundance
Untreated
B
Shannon index
Uninvaded
Ec040
Ec069
Ec131
Ec744
C
With Antibiotic
D

Figure S6. Microbial community composition and relative abundance at the end of the 'live' microcosm assay (see Fig. 1) in untreated microcosms (A) & (B) and in antibiotic-treated microcosms (C) & (D). (A) Relative abundance of genera present in antibiotic-free microcosms after 48 hours with and without introduced focal strains for each human donor (Donor1, Donor2, Donor3). (C) Relative abundance of genera present in antibiotic-treated microcosms with and without introduced focal strains after 48 hours for each human gut microbiome (Donor1, Donor2, Donor3). We detected an average of 26.87 ASVs in antibiotic-treated microcosms, compared with 34.37 ASVs in microcosms without antibiotics (ANOVA: antibiotic effect - $F(1,60)= 295.652$, $p<0.001$; donor effect - $F(2,60)= 39.056$, $p<0.001$; antibiotic-donor interaction - $F(2,60)= 19.813$, $p<0.001$). (B) & (D) Shannon's diversity index for microcosms in each treatment group. On average Shannon's diversity index was lower in microcosms with vs without antibiotics (ANOVA: antibiotic effect - $F(1,60)= 56.213$, $p<0.001$; donor effect - $F(2,60)= 148.281$, $p<0.001$; antibiotic-donor interaction - $F(2,60)= 45.343$, $p<0.001$; strain-donor interaction: $F(8,60)= 2.192$, $p<0.05$; antibiotic-strain-donor interaction: $F(8,60)= 2.198$, $p<0.05$). An increase in the Shannon index can result from higher species evenness, even if species richness remains unchanged. In the case of Donor2, the observed increase in the Shannon index appears to be driven by higher evenness under antibiotic treatment (see also Figures S4 and S7).

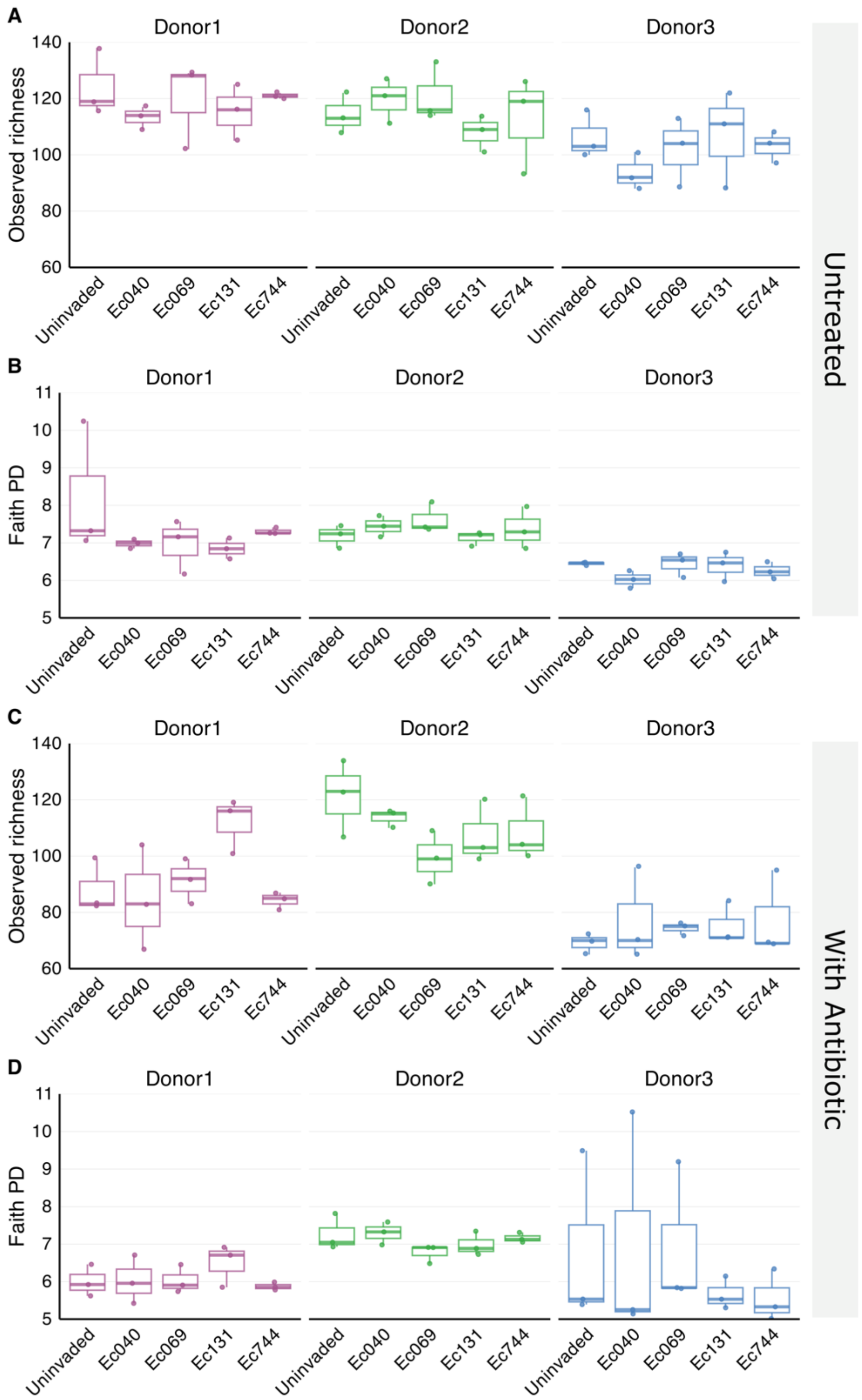


Figure S7. Alternative measures of alpha diversity, observed richness and Faith phylogenetic diversity (PD), shown for both antibiotic-free and antibiotic-treated groups of microcosms.

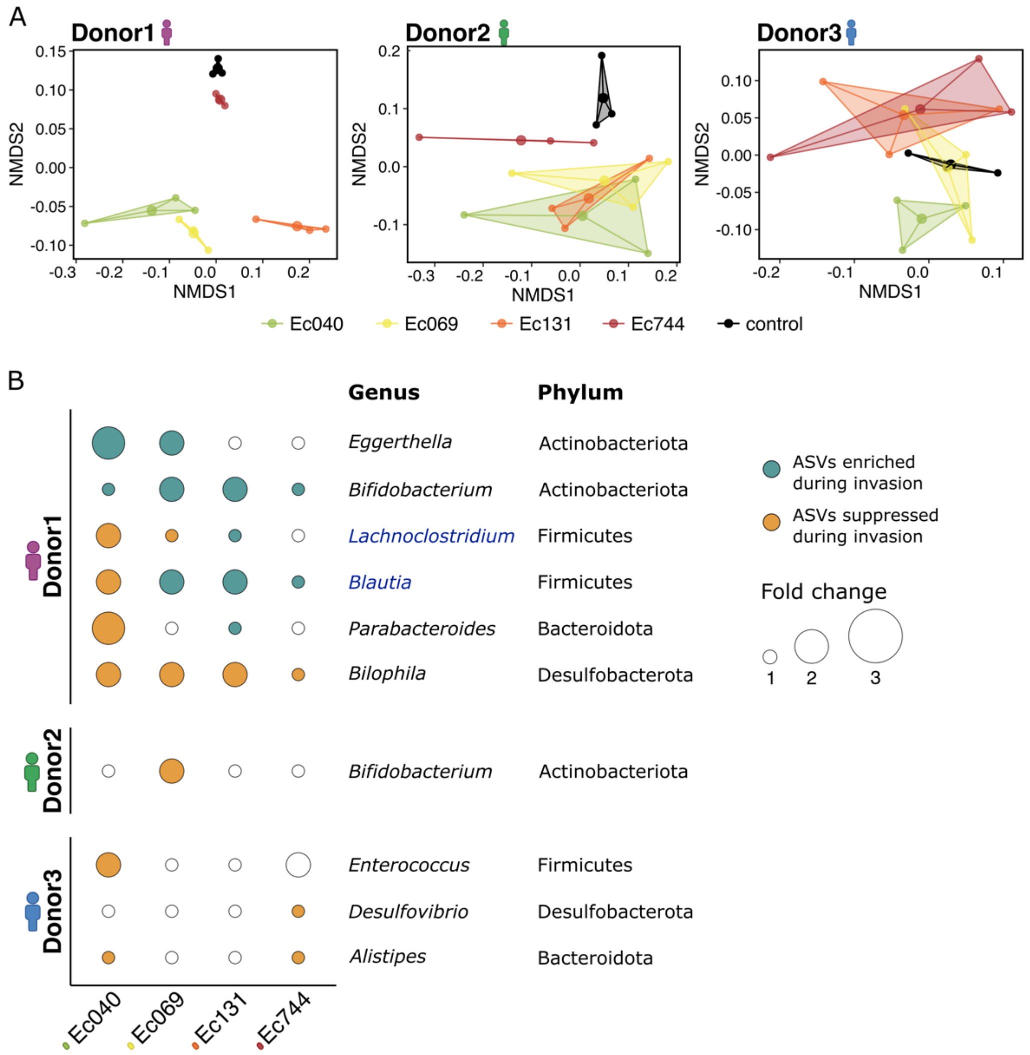


Figure S8. Strain-specific interactions between incoming resistant *E. coli* and resident microbiota in antibiotic-treated gut microcosms. (A) Non-metric multidimensional scaling (nMDS) of microbial community composition (genus level based on 16S rRNA) from Bray-Curtis distance matrices, showing all microcosms in groups inoculated with different resistant strains, including an uninoculated control (see legend at bottom), with antibiotic treatment for each human donor (panels). Final community composition varied depending on the identity of the invading strain (strain effect in PERMANOVA, excluding uninvaded microcosms: $F(3,24)= 2.766$, $p<0.01$), and the sample donor (donor effect: $F(2,24)= 107.826$, $p<0.001$). Differences among strains depended on the donor (strain × donor interaction: $F(6,24)= 2.220$, $p<0.05$), with the greatest differences observed for Donor1 (strain effect tested within each donor: $p<0.001$ for Donor1, $p>0.05$ for Donor2 and Donor3). The *Escherichia* genus is excluded from these analyses. (B) Linear Discriminant Analysis Effect Size (LEfSe) results showing the amplicon sequence variants (ASVs) enriched or suppressed in microcosms inoculated with each of the focal strains vs uninoculated microcosm groups (rows), for the set of microcosms with each incoming resistant strain (columns) and from each human donor (groups of rows). Filled circles show combinations with a LDA (Linear discriminant analysis) score >2; circle colour shows the direction of the effect (enriched/ suppressed) and circle size indicates effect size (fold change, see legend). Only taxa showing a significant difference between inoculated and uninoculated microcosms ($p < 0.05$) in at least one combination are shown. Genera highlighted in blue belong to the *Lachnospiraceae* family.

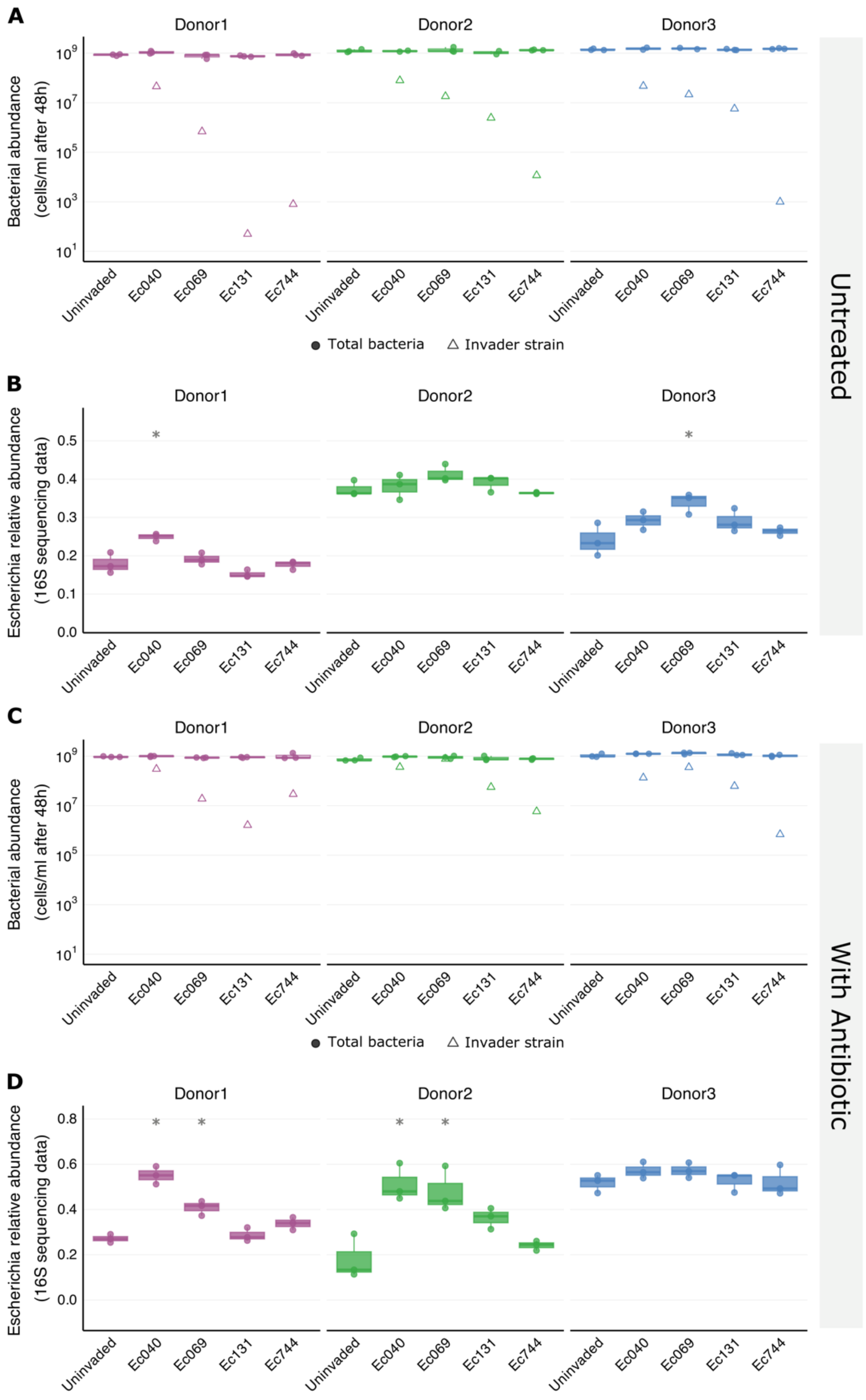


Figure S9. (A) & (C) Total bacterial abundance at the end of the ‘live’ microcosm assay (48 hours; measured by flow cytometry, see Methods); the mean abundance of incoming strains is also shown for reference (triangles - measured by colony counting). In all treatments with incoming strains, the total bacterial abundance was similar to that in the equivalent uninoculated control microcosms ($p>0.05$ in all cases, tested by pairwise $t$-testing with sequential Bonferroni). (B & D) Relative abundance of *Escherichia* detected by 16S rRNA gene sequencing at the end of the main experiment (see also Figure S6). In some but not all groups, *Escherichia* relative abundance was higher in inoculated than in equivalent uninoculated microcosms (asterisks show groups where $p<0.05$, tested by pairwise *t*-testing with sequential Bonferroni correction).

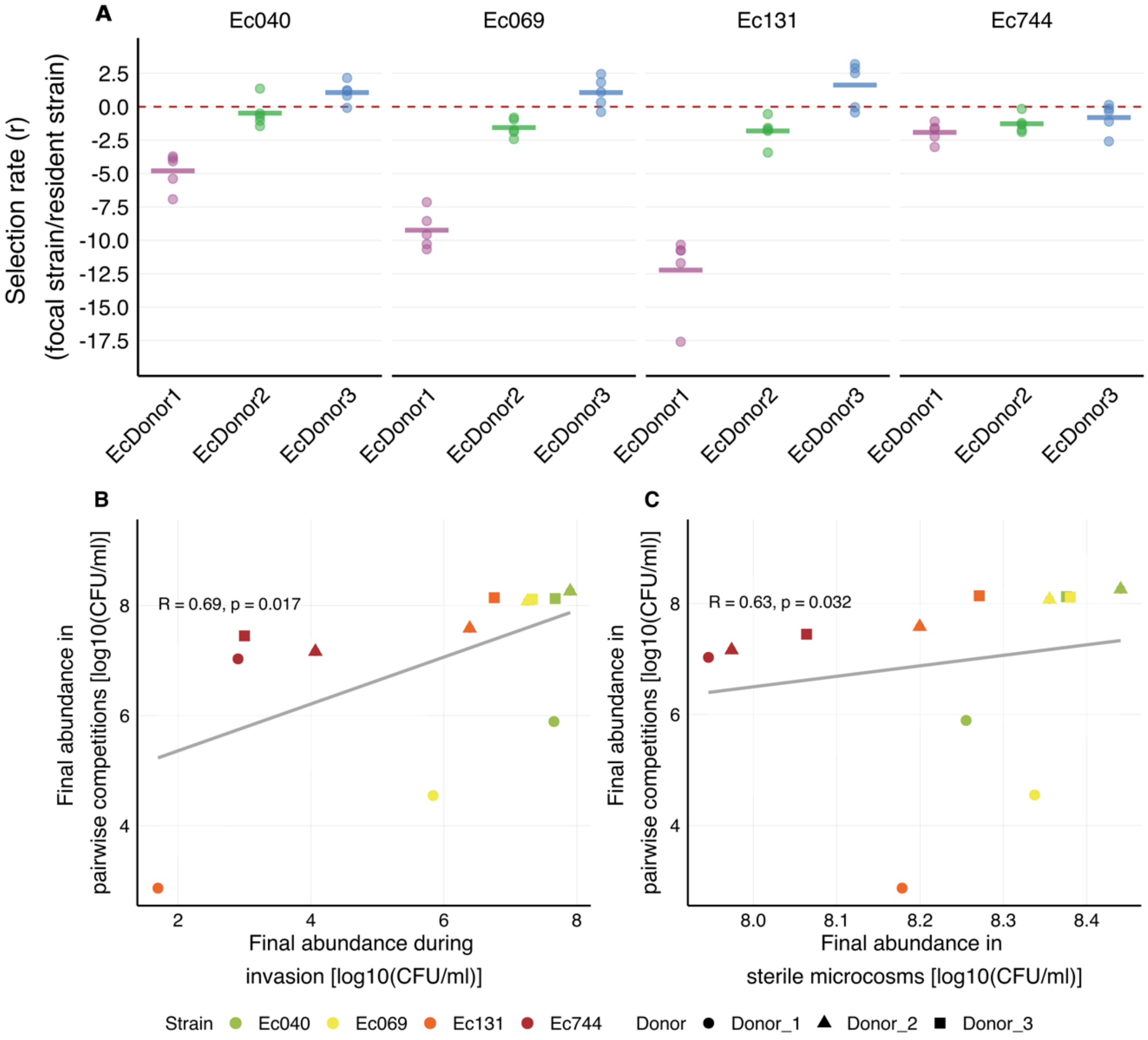


Figure S10. Competitive interactions between incoming resistant *E. coli* and resident *E. coli*. (A) Selection rate from pairwise competitions between the four incoming strains and the three *E. coli* resident strains (EcDonor1, EcDonor2, EcDonor3) (corresponds to Fig. 4 in the main text). (B) Positive correlation between the final abundance of each incoming strain in the pairwise competitions with resident *E. coli* and the incoming strain final abundance during the 'live' microcosm assay (corresponding to Fig. 4 and Fig. 1, respectively). (C) Positive correlation between final abundance of each incoming strain in pairwise competitions with resident *E. coli* and the incoming strain final abundance in sterilised microcosms (corresponding to Fig. 4 and Fig. 2, respectively). In (B)&(C), colours represent the incoming strains, shapes represent the donor sample (see legend at bottom).

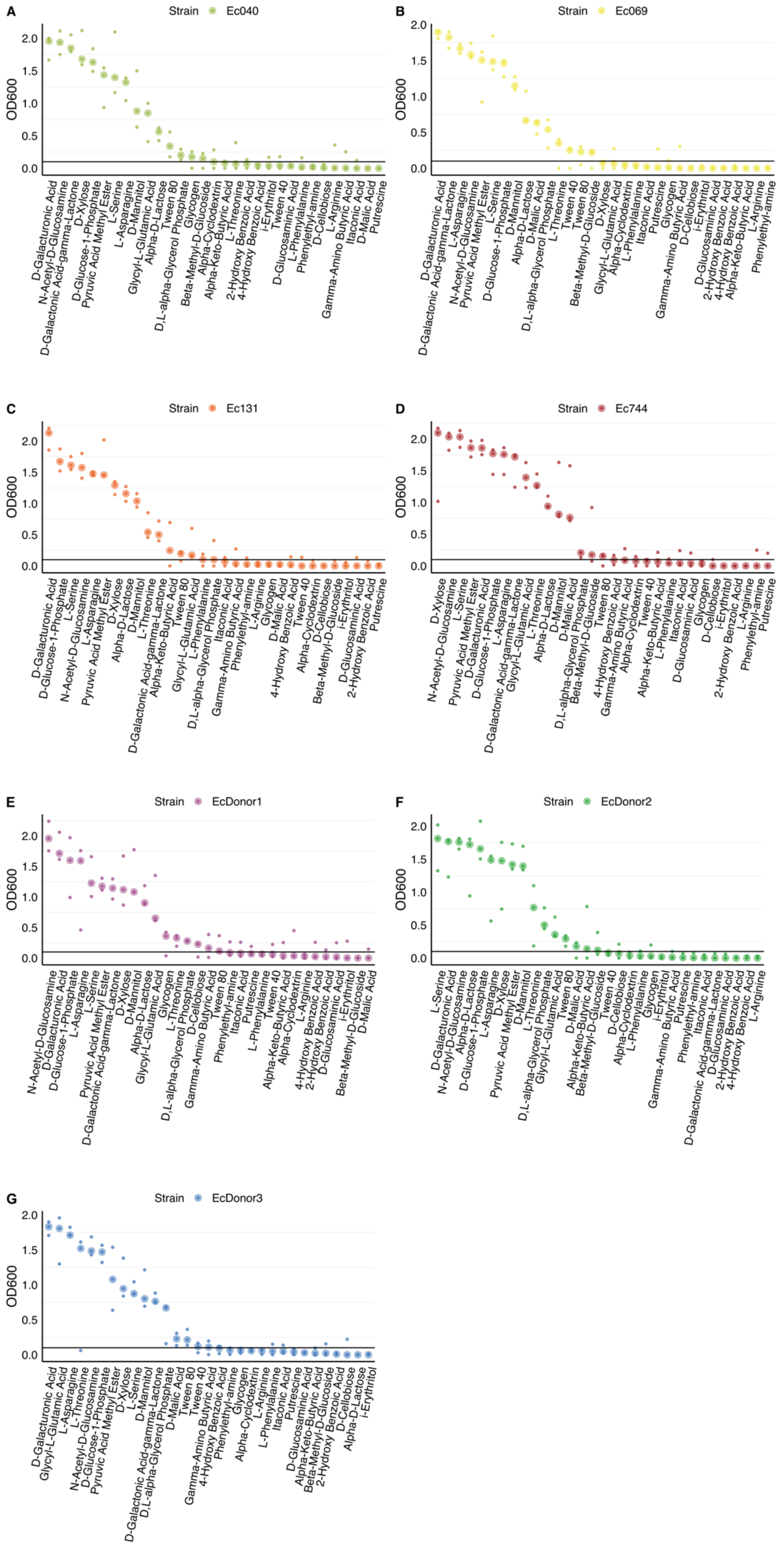

A
Strain Ec040
OD600
0.0 0.5 1.0 1.5 2.0
D-Galacturonic Acid
N-Acetyl-D-Glucosamine
D-Galactonic Acid-gamma-Lactone
D-Xylose
D-Glucose-1-Phosphate
Pyruvic Acid Methyl Ester
L-Serine
L-Asparagine
D-Mannitol
Glycyl-L-Glutamic Acid
Alpha-D-Lactose
Tween 80
D,L-alpha-Glycerol Phosphate
Glycogen
Beta-Methyl-D-Glucoside
Alpha-Cyclodextrin
Alpha-Keto-Butyric Acid
L-Threonine
2-Hydroxy Benzoic Acid
4-Hydroxy Benzoic Acid
i-Erythritol
Tween 40
D-Glucosaminic Acid
L-Phenylalanine
Phenylethyl-amine
D-Cellobiose
Gamma-Amino Butyric Acid
L-Arginine
Itaconic Acid
D-Malic Acid
Putrescine
B
Strain Ec069
OD600
0.0 0.5 1.0 1.5 2.0
D-Galacturonic Acid
D-Galactonic Acid-gamma-Lactone
L-Asparagine
N-Acetyl-D-Glucosamine
Pyruvic Acid Methyl Ester
L-Serine
D-Glucose-1-Phosphate
D-Mannitol
Alpha-D-Lactose
D-Malic Acid
D,L-alpha-Glycerol Phosphate
L-Threonine
Tween 40
Tween 80
Beta-Methyl-D-Glucoside
D-Xylose
Glycyl-L-Glutamic Acid
Alpha-Cyclodextrin
L-Phenylalanine
Itaconic Acid
Putrescine
Gamma-Amino Butyric Acid
Glycogen
D-Cellobiose
i-Erythritol
D-Glucosaminic Acid
2-Hydroxy Benzoic Acid
4-Hydroxy Benzoic Acid
Alpha-Keto-Butyric Acid
L-Arginine
Phenylethyl-amine
C
Strain Ec131
OD600
0.0 0.5 1.0 1.5 2.0
D-Galacturonic Acid
D-Glucose-1-Phosphate
L-Serine
N-Acetyl-D-Glucosamine
L-Asparagine
Pyruvic Acid Methyl Ester
D-Xylose
Alpha-D-Lactose
D-Mannitol
D-Galactonic Acid-gamma-Lactone
L-Threonine
Alpha-Keto-Butyric Acid
Tween 80
Glycyl-L-Glutamic Acid
L-Phenylalanine
D,L-alpha-Glycerol Phosphate
Itaconic Acid
Gamma-Amino Butyric Acid
Phenylethyl-amine
L-Arginine
Glycogen
4-Hydroxy Benzoic Acid
D-Malic Acid
Alpha-Cyclodextrin
Tween 40
Beta-Methyl-D-Glucoside
D-Cellobiose
i-Erythritol
D-Glucosaminic Acid
2-Hydroxy Benzoic Acid
Putrescine
D
Strain Ec744
OD600
0.0 0.5 1.0 1.5 2.0
D-Xylose
N-Acetyl-D-Glucosamine
L-Serine
Pyruvic Acid Methyl Ester
D-Galacturonic Acid
D-Glucose-1-Phosphate
D-Galactonic Acid-gamma-Lactone
L-Asparagine
Glycyl-L-Glutamic Acid
L-Threonine
Alpha-D-Lactose
D-Mannitol
D-Malic Acid
D,L-alpha-Glycerol Phosphate
Beta-Methyl-D-Glucoside
Tween 80
4-Hydroxy Benzoic Acid
Gamma-Amino Butyric Acid
Alpha-Cyclodextrin
Tween 40
Alpha-Keto-Butyric Acid
L-Phenylalanine
Itaconic Acid
D-Glucosaminic Acid
Glycogen
D-Cellobiose
i-Erythritol
2-Hydroxy Benzoic Acid
L-Arginine
Phenylethyl-amine
Putrescine
E
Strain EcDonor1
OD600
0.0 0.5 1.0 1.5 2.0
N-Acetyl-D-Glucosamine
D-Galacturonic Acid
D-Glucose-1-Phosphate
L-Asparagine
L-Serine
Pyruvic Acid Methyl Ester
D-Galactonic Acid-gamma-Lactone
D-Xylose
D-Mannitol
Alpha-D-Lactose
Glycyl-L-Glutamic Acid
Glycogen
D,L-alpha-Glycerol Phosphate
L-Threonine
D-Cellobiose
Gamma-Amino Butyric Acid
Tween 80
Phenylethyl-amine
Itaconic Acid
Putrescine
L-Phenylalanine
Tween 40
Alpha-Keto-Butyric Acid
Alpha-Cyclodextrin
L-Arginine
4-Hydroxy Benzoic Acid
2-Hydroxy Benzoic Acid
D-Glucosaminic Acid
i-Erythritol
Beta-Methyl-D-Glucoside
D-Malic Acid
F
Strain EcDonor2
OD600
0.0 0.5 1.0 1.5 2.0
L-Serine
D-Galacturonic Acid
N-Acetyl-D-Glucosamine
Alpha-D-Lactose
D-Glucose-1-Phosphate
L-Asparagine
D-Xylose
Pyruvic Acid Methyl Ester
D-Mannitol
L-Threonine
D,L-alpha-Glycerol Phosphate
Glycyl-L-Glutamic Acid
Tween 80
D-Malic Acid
Alpha-Keto-Butyric Acid
Beta-Methyl-D-Glucoside
Tween 40
D-Cellobiose
Alpha-Cyclodextrin
L-Phenylalanine
Glycogen
i-Erythritol
Gamma-Amino Butyric Acid
Putrescine
Phenylethyl-amine
Itaconic Acid
D-Galactonic Acid-gamma-Lactone
D-Glucosaminic Acid
2-Hydroxy Benzoic Acid
4-Hydroxy Benzoic Acid
L-Arginine
G
Strain EcDonor3
OD600
0.0 0.5 1.0 1.5 2.0
D-Galacturonic Acid
Glycyl-L-Glutamic Acid
L-Asparagine
L-Threonine
N-Acetyl-D-Glucosamine
D-Glucose-1-Phosphate
Pyruvic Acid Methyl Ester
D-Xylose
L-Serine
D-Mannitol
D-Galactonic Acid-gamma-Lactone
D,L-alpha-Glycerol Phosphate
D-Malic Acid
Tween 80
Tween 40
Gamma-Amino Butyric Acid
4-Hydroxy Benzoic Acid
Phenylethyl-amine
Glycogen
Alpha-Cyclodextrin
L-Arginine
L-Phenylalanine
Itaconic Acid
Putrescine
D-Glucosaminic Acid
Alpha-Keto-Butyric Acid
Beta-Methyl-D-Glucoside
2-Hydroxy Benzoic Acid
D-Cellobiose
Alpha-D-Lactose
i-Erythritol

Figure S11. (A-G). Bacterial growth (OD600) shown for each carbon source in the BioLog EcoPlates for the four incoming strains and the three resident *E. coli* strains. Small points show replicates and larger points show the average in each combination. The horizontal line indicates a cut-off value to assign carbon sources as being used (positive impact on growth) or not by a given strain; we defined the cut-off as median OD590 nm after subtracting the blank > 0.1 (see also Figure 5 in the main text).

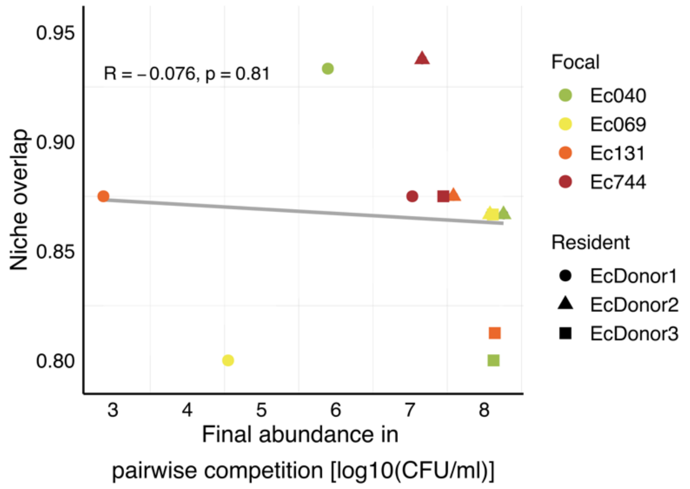


Figure S12. Scatter plot between the final growth of each incoming strain in the pairwise competitions against the resident *E. coli* strains (x-axis, see also Fig. 4 in the main text) and their niche overlap with each of the three resident strains (y-axis, see also Fig. 5E in the main text) showing no significant correlation.

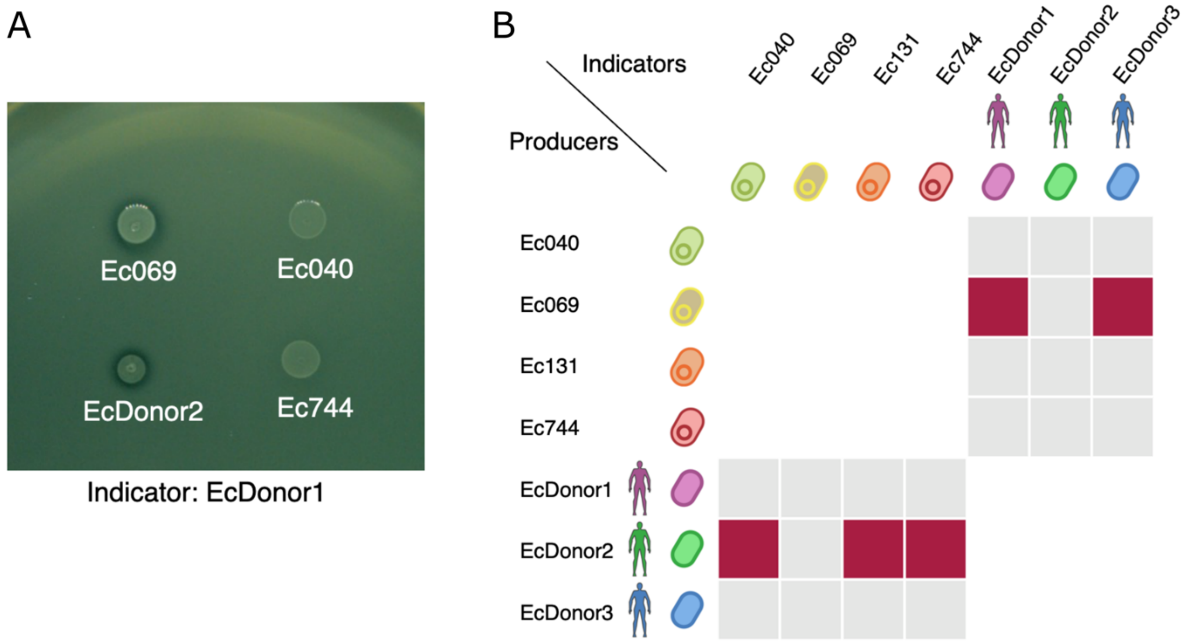


Figure S13. Inhibitory interactions among incoming resistant strains and resident *E. coli* strains. (A) Representative images from agar overlay assays. 'Producers', labelled in the image, were spotted onto a lawn of an 'indicator' strain. (B) Results of inhibition assays testing whether each resistant incoming strain could inhibit each resident *E. coli* strain (as producer and indicator, respectively), and vice versa. Red squares indicate combinations where a halo was visible in all three replicate assays (growth inhibition of the indicator by the producer).

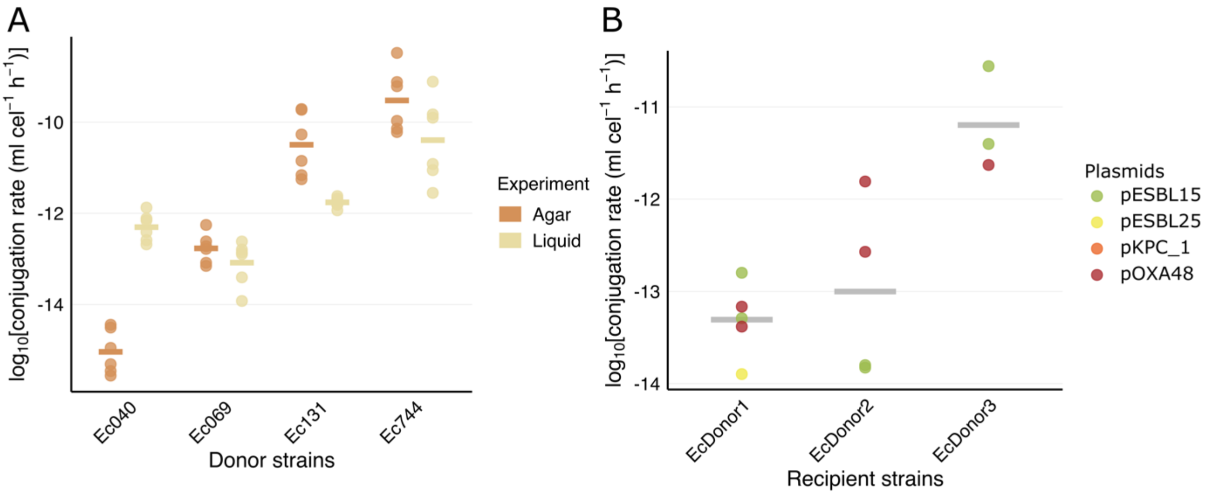


Figure S14. (A) Conjugation rate for each of the four plasmids (see Table S2) from their original clinical host strain (incoming strains) to *E. coli* MG1655. Each combination was tested both in liquid and on agar (legend at right). For these assays, bacterial donors and recipients were streaked from freezer stocks onto solid LB agar with ampicillin 100 µg/ml and chloramphenicol 25 µg/ml, respectively, and incubated overnight at 37 °C. Donor and recipient colonies were independently inoculated in 2 ml LB and incubated overnight. After growth, donor and recipient cultures were collected by centrifugation (15 min, 1500 × g) and cells were re-suspended with 300 µl sterile NaCl 0.9%. Then, suspensions were mixed 1:1 *v*:*v*, spotted onto solid LB medium and incubated 37 °C one hour or in liquid LB overnight. Transconjugants were selected by streaking the conjugation mix on LB with ampicillin 100 µg/ml and chloramphenicol 25 µg/ml, donors+transconjugants with ampicillin 100 µg/ml, and recipients+transconjugants with chloramphenicol 25 µg/ml. Conjugation rates were determined using the end-point method[1]. (B) Conjugation rate for the four plasmids (legend) from a diaminopimelic acid (DAP) auxotrophic laboratory mutant of *E. coli* K-12, transferring to the three *E. coli* resident strains (*x*-axis) in filter matting assays with LB agar. Here, to be able to select for transconjugants, we first performed an initial conjugation round to introduce each focal plasmid into *E. coli* β3914, a diaminopimelic acid (DAP) auxotrophic laboratory mutant of *E. coli* K-12 (kanamycin, erythromycin, and tetracycline resistant). Then, we performed a conjugation experiment as above, using the DAP transconjugants as secondary donors and the resident *E. coli* strains as recipients. Transconjugants were selected by streaking the conjugation mix on LB with ampicillin 100 µg/ml (the donors are counter-selected without DAP), donors+transconjugants with ampicillin 100 µg/ml and DAP 0.3mM, and recipients+transconjugants with LB agar (the donors are counter-selected without DAP).

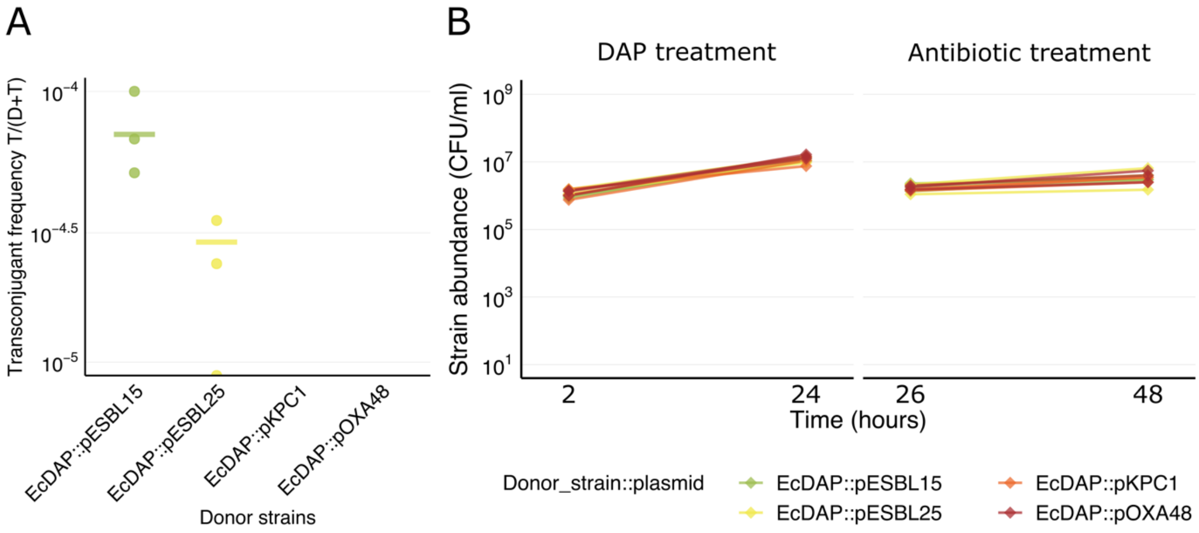


Figure S15. (A) Transconjugant frequency after 48h in anaerobic 'live' gut microcosms prepared with the Donor1 microbiome sample, and DAP *E. coli* β3914 as the donor strain with each plasmid (*x*-axis). Points show replicates; no points are shown in combinations where no transconjugants were detected. Plating here was on LB with ampicillin 100 µg/ml with/ without DAP 0.3mM, to count both the total abundance and the transconjugants (counter-selecting the donors). Final transconjugant frequencies are shown as transconjugants/(donors+transconjugants) (T/(D+T)). (B) Abundance of the DAP *E. coli* K12 strain with each plasmid (legend) during the conjugation experiment in the anaerobic microcosms prepared with the Donor1 sample. DAP was applied during the first 24 hours, followed by transfer to fresh microcosms without DAP but with ampicillin after 24 hours. Abundance of the DAP *E. coli* β3914 strain was independent of which plasmid it carried initially, both after 24 h (one-way ANOVA: $F(3,8)= 1.56$, $p>0.05$) and after 48 h (one-way ANOVA: $F(3,8)= 0.25$, $p>0.05$).

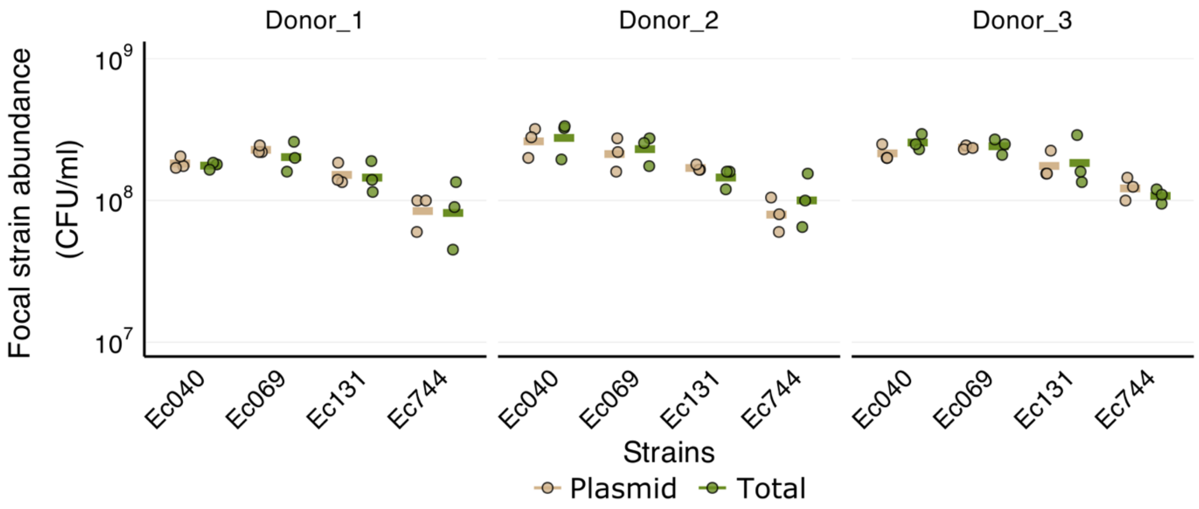


Fig S16. Plasmid stability in gut microcosms under our experimental conditions (sterilised microcosms for 48h). Abundance of each strain estimated by colony counts on agar plates selecting for plasmid-encoded resistance phenotypes ("Plasmid") and on non-selective plates ("Total") after incubation without antibiotics in our experimental conditions (sterilised microcosms for 48h). For each donor sample (panels, left to right), abundance of each resistant focal strain (*x*-axis) is shown after 48 h incubation in sterilised, anaerobic, antibiotic-free microcosms prepared as in the main experiment. Samples from three replicate microcosms in each combination were plated on agar selecting for resistance phenotypes encoded by the relevant plasmids (chromID ESBL and CARBA SMART agar; bioMérieux, Switzerland) and on non-selective plates (chromatic agar without antibiotics; Chromatic MH, Liofilchem, Roseto degli Abruzzi, Italy). The extremely similar counts on the two types of plates in all combinations indicate plasmids were retained at stable frequencies during the experiment.

Table S1. Bacterial strains used in this study. ST=Sequence Type.

| **Strain_ID** | **Alternative_ID** | **Species** | **ST** | **Phylogroup** | **Plasmid_replicons** | **Comment** | **Reference** |
|---|---|---|---|---|---|---|---|
| **Ec040** | **ESBL15** | *E. coli* | 40 | B1 | IncI1, ColRNAI, Col156 | Clinical antibiotic-resistant *E. coli* strain (focal strain) | Benz *et al*., 2020; Tschudin-Sutter *et al*., 2016 |
| **Ec069** | **ESBL25** | *E. coli* | 69 | D | IncFIB, IncFIA, IncFII, p0111, IncB, ColRNAI, Col156, ColRNAI, Col8282, Col(MG828) | Clinical antibiotic-resistant *E. coli* strain (focal strain) | Benz *et al*., 2020; Tschudin-Sutter *et al*., 2016 |
| **Ec131** | **KPC_1** | *E. coli* | 131 | B2 | IncFIA, IncFII, IncFIA, IncL, IncX3, IncU, IncX3, Col8282 | Clinical antibiotic-resistant *E. coli* strain (focal strain) | Noll, *et al*., 2018. |
| **Ec744** | **Oxa48_1** | *E. coli* | 744 | A | IncFIC, IncFII, IncFIB, IncL, ColRNAI, Col(MG828) | Clinical antibiotic-resistant *E. coli* strain (focal strain) | Noll, *et al*., 2018. |
| **EcDonor1** | **NI2** | *E. coli* | 95 | B2 | IncB, IncX3 | Most abundant *E. coli* Isolated from human sample Donor1 | Boumasmoud *et al*. 2024 |
| **EcDonor2** | **NI3** | *E. coli* | ~10 | A | IncFIB | Most abundant *E. coli* Isolated from human sample Donor2 | Boumasmoud *et al*. 2024 |
| **EcDonor3** | **NI4** | *E. coli* | 1193 | B2 | IncFIA | Most abundant *E. coli* Isolated from human sample Donor3 | Boumasmoud *et al*. 2024 |
| **EcDAP** | **β3914** | *E. coli* | | A | - | Diaminopimelic acid (DAP) auxotrophic laboratory mutant of *E. coli* K-12 | Alonso-del-Valle *et al*., 2021 |

Table S2. ESBL and carbapenemase plasmids carried by focal resistant *E. coli* strains

| **Plasmid** | **Bacterial_host** | **Size (kb)** | **Incompatibility group** | **Resistance_genes** | **Reference (Genbank Ac. No.)** |
|---|---|---|---|---|---|
| pESBL15 | *E. coli* Ec040 | 88.9 | IncI | $bla_{CTX\text{-}M\text{-}1}$ | SAMN12275742 |
| pESBL25 | *E. coli* Ec069 | 131 | IncFIB, IncFIA | $bla_{CTX\text{-}M\text{-}14}$, *tetB*, *dfrA*, *mphA* | SAMN17073781 |
| pKPC | *E. coli* Ec131 | 73.7 | IncX3, IncU | $bla_{KPC2}$, *sat2A* | UWWZ01000004.1 |
| pOXA48 | *E. coli* Ec744 | 63.6 | IncL | $bla_{OXA\text{-}48}$ | UWXP01000003.1 |

Table S3. Primers used for plasmid identification and strain identification (REP-PCR).

| **Name/ Target plasmid** | **Target gene** | **Forward primer 5' - 3'** | **Reverse primer 5' - 3'** |
|---|---|---|---|
| pESBL15 | CTXM-55 | CCGCGGTGCTGAAGAAAAGT | TCATCCATGTCACCAGCTGC |
| pESBL25 | CTXM-14 | GCGGCTGGGTAAAATAGGTC | GAGAGTGCAACGGATGATGT |
| pKPC | KPC | TGTGCAGCTCATTCAAGGGC | GCCTCGCTGTGCTTGTCATC |
| pOXA-48 | OXA-48 | GGCGTAGTTGTGCTCTGGAA | CCAACCGACCCACCAGCCAA |
| REP-PCR | - | NNNGCGCCGNCATCAGGC | ACGTCTTATCAGGCCTAC |
